# scRNASeqDB: a database for gene expression profiling in human single cell by RNA-seq

**DOI:** 10.1101/104810

**Authors:** Yuan Cao, Junjie Zhu, Guangchun Han, Peilin Jia, Zhongming Zhao

## Abstract

Summary: Single-cell RNA sequencing (scRNA-Seq) is quickly becoming a powerful tool for high-throughput transcriptomic analysis of cell states and dynamics. Both the number and quality of scRNA-Seq datasets have dramatically increased recently. So far, there is no database that comprehensively collects and curates scRNA-Seq data in humans. Here, we present scRNASeqDB, a database that includes almost all the currently available human single cell transcriptome datasets (n= 36) covering 71 human cell lines or types and 8910 samples. Our online web interface allows user to query and visualize expression profiles of the gene(s) of interest, search for genes that are expressed in different cell types or groups, or retrieve differentially expressed genes between cell types or groups. The scRNASeqDB is a valuable resource for single cell transcriptional studies.

Availability: The database is available at https://bioinfo.uth.edu/scrnaseqdb/.

## 1 Introduction

Single-cell RNA sequencing (scRNA-Seq) is rapidly evolving as a powerful tool for high-throughput transcriptomic analysis of cell states and dynamics. It can accurately measure RNA expression characteristics at single cell level in order to explore cell phenotype, function and transcriptomic heterogeneity (Linnarsson and Teichmann, 2016), which is averaged in typical bulk tissue based studies (Shalek et al., 2013). During the past a few years, both the quality and quantity of scRNA-Seq data have dramatically increased, calling for a centralized web resource that curates and provides features of the single-cell gene expressing profiles in broad research community. So far, there have been three small-scale web resources for single cell transcriptome data, all of which were developed for mouse data (Du et al., 2015; Nestorowa et al., 2016; Biase et al., 2014). Du et al. (2015) developed a web resource for only one single-cell gene expression dataset in the developing mouse lung tissue. Nestorowa et al. (2016) built a web interface using only one HPSC transcriptome dataset, providing insights into differentiation of blood stem cell in mice. And Biase et al. (2014) developed an online genome browser for mouse scRNA-Seq data. To our knowledge, there has been no database for curation and annotation of human scRNA-Seq data yet. Furthermore, the above databases were mainly based on one project or dataset.

Here, we present scRNASeqDB, a database that collects and curates all the publicly available human single cell gene expression datasets. Through metadata annotation and unified data processing procedures, it covers 71 human cell lines or types and 8910 samples. The data and follow-up analysis results are available through an intuitive web interface. This timely database will facilitate researchers for gene expression studies in human single cells in broad biology and medicine fields.

## 2 Design and implementation

### 2.1 Data collection

We searched the NCBI Gene Expression Omnibus (GEO) database (Clough and Barrett, 2016) for gene expression profiling experiments with the keywords such as scRNAseq, single-cell RNA-Seq, single-cell transcriptome. A detailed description is provided in Table S1. We carefully reviewed the resultant papers and datasets. In total, we obtained 36 datasets for human single cell RNA-Seq analysis (Table S2). These datasets included 71 human cell lines or types, which are related to reproduction, immune system, the brain and nervous system, cancer, stem cell, etc. (Table S3). The meta data of GEO datasets was downloaded and imported into MySQL by using the R package GEOquery (Davis and Meltzer, 2007) and RMYSQL, respectively. For RNA-Seq experiments, the gene expression matrices were also retrieved from the GEO and convert to transcripts per kilobase million (TPM) or read count format by using our in-house R scripts. Cell samples were manually grouped according to the characteristics of the dataset or description in the original publications (Fig S1).

### 2.2 Web interface

Currently, scRNASeqDB release 1.0 has a collection of 8910 samples belonging to 174 cell groups from 36 datasets. The web interface of scRNASeqDB was implemented in PHP and JavaScript by using Yii framework, which enables user to search across the database easily without requirement of computer expertise. Interactive heatmap and box plots are constructed dynamically to display gene expression for individual cells and cell groups by using the HighCharts component. Qtlcharts (Broman, 2015) is used to present correlation matrices and scatter plots.

### 2.3 Database features

The main purpose of scRNASeqDB is to facilitate analysis and visualization of gene expression profiles across various human single cells. The ‘Gene View’ page displays the expression profile of a gene (Fig S2). It consists of three sections. (1) The general information section includes gene symbol, Ensembl ID, description, etc. To enable user to further explore gene information, it provides web links of the gene to other resources, such as OMIM (Amberger et al., 2009) and Ensembl (Aken et al., 2016). (2) In the gene expression section, an interactive heatmap is available to display gene expression across individual cells in each dataset. A box plot and a table summarizing the minimum, median, and maximum value of the gene expression in each cell group from each dataset are also provided. If there are more than one cell group in a dataset, comparison of query gene expression between cell groups will be listed to indicate significantly expressed gene. (3) It also shows the top 100 positively and negatively correlated genes across all cells in a dataset for the query gene. GO and KEGG pathway annotations (Huang et al., 2009) for these genes are provided too. In the ‘Cell View’ page, user can explore the top regulated genes in the query cell group and the relationship between two genes in the cell group (Fig S3). Through ‘Dataset View’, user can obtain comprehensive description and cell groups in the query dataset. Meanwhile, lists of differentially expressed genes between two cell groups in the dataset are displayed, as well as GO and KEGG pathway enrichment results for the differentially expressed genes (Fig S4).

## 3 Conclusions

scRNASeqDB is a user-friendly database that timely collects and curates gene expression profiles of the transcriptome of human single cells. The database currently has 36 datasets covering the gene expression of 8910 single cells from 174 cell groups. It provides various features such as gene expression in different cell types, expression patterns at the pathway and network levels, and tools for visualization and exploration of gene expression in single cells. In comparison with other resources, scRNASeqDB is by far the most comprehensive database for curated and analyzed scRNA-Seq data and the only one for human scRNA-Seq data. It will be routinely maintained to include future scRNA-Seq data.

## Acknowledgements

None

## References

Aken, B.L. et al. (2016) The Ensembl gene annotation system. Database, 2016, baw093.

Amberger, J. et al. (2009) McKusick’s Online Mendelian Inheritance in Man (OMIM). Nucleic Acids Res., 37, D793–796.

Biase, F.H. et al. (2014) Cell fate inclination within 2-cell and 4-cell mouse embryos revealed by single-cell RNA sequencing. Genome Res., 24, 1787–1796.

Broman, K.W. (2015) R/qtlcharts: interactive graphics for quantitative trait locus mapping. Genetics, 199, 359–361.

Clough, E. and Barrett, T. (2016) The Gene Expression Omnibus Database. Methods Mol. Biol., 1418, 93–110.

Davis, S. and Meltzer, P.S. (2007) GEOquery: a bridge between the Gene Expression Omnibus (GEO) and BioConductor. Bioinformatics, 23, 1846–1847.

Du, Y. et al. (2015) ‘LungGENS’: a web-based tool for mapping single-cell gene expression in the developing lung. Thorax, 70, 1092–1094.

Huang, D.W. et al. (2009) Systematic and integrative analysis of large gene lists using DAVID bioinformatics resources. Nat. Protoc., 4, 44–57.

Linnarsson, S. and Teichmann, S.A. (2016) Single-cell genomics: coming of age. Genome Biol., 17, 97.

Nestorowa, S. et al. (2016) A single-cell resolution map of mouse hematopoietic stem and progenitor cell differentiation. Blood, 128, e20–31.

Shalek, A.K. et al. (2013) Single-cell transcriptomics reveals bimodality in expression and splicing in immune cells. Nature, 498, 236–240.

